# Elucidating the influence of RNA modifications and Magnesium ions on tRNA^Phe^ conformational dynamics in *S. cerevisiae*: Insights from Replica Exchange Molecular Dynamics simulations

**DOI:** 10.1101/2024.03.11.584441

**Authors:** Zohreh R. Nowzari, Rebecca J D’Esposito, Sweta Vangaveti, Alan A. Chen

## Abstract

Post-transcriptional modifications in RNA can significantly impact their structure and function. In particular, transfer RNAs (tRNAs) are heavily modified, with around 100 different naturally occurring nucleotide modifications contributing to codon bias and decoding efficiency. Here, we describe our efforts to investigate the impact of RNA modifications on the structure and stability of tRNA Phenylalanine (tRNA^Phe^) from S. *cerevisiae* using molecular dynamics (MD) simulations. Through temperature replica exchange MD (T-REMD) studies, we explored the unfolding pathway to understand how RNA modifications influence the conformational dynamics of tRNA^Phe^, both in the presence and absence of magnesium ions (Mg^2+^). We observe that modified nucleotides in key regions of the tRNA establish a complex network of hydrogen bonds and stacking interactions which is essential for tertiary structure stability of the tRNA. Furthermore, our simulations show that modifications facilitate the formation of ion binding sites on the tRNA. However, high concentrations of Mg^2+^ ions can stabilize the tRNA tertiary structure in the absence of modifications. Our findings illuminate the intricate interactions between modifications, magnesium ions, and RNA structural stability.

**Graphical Abstract:** 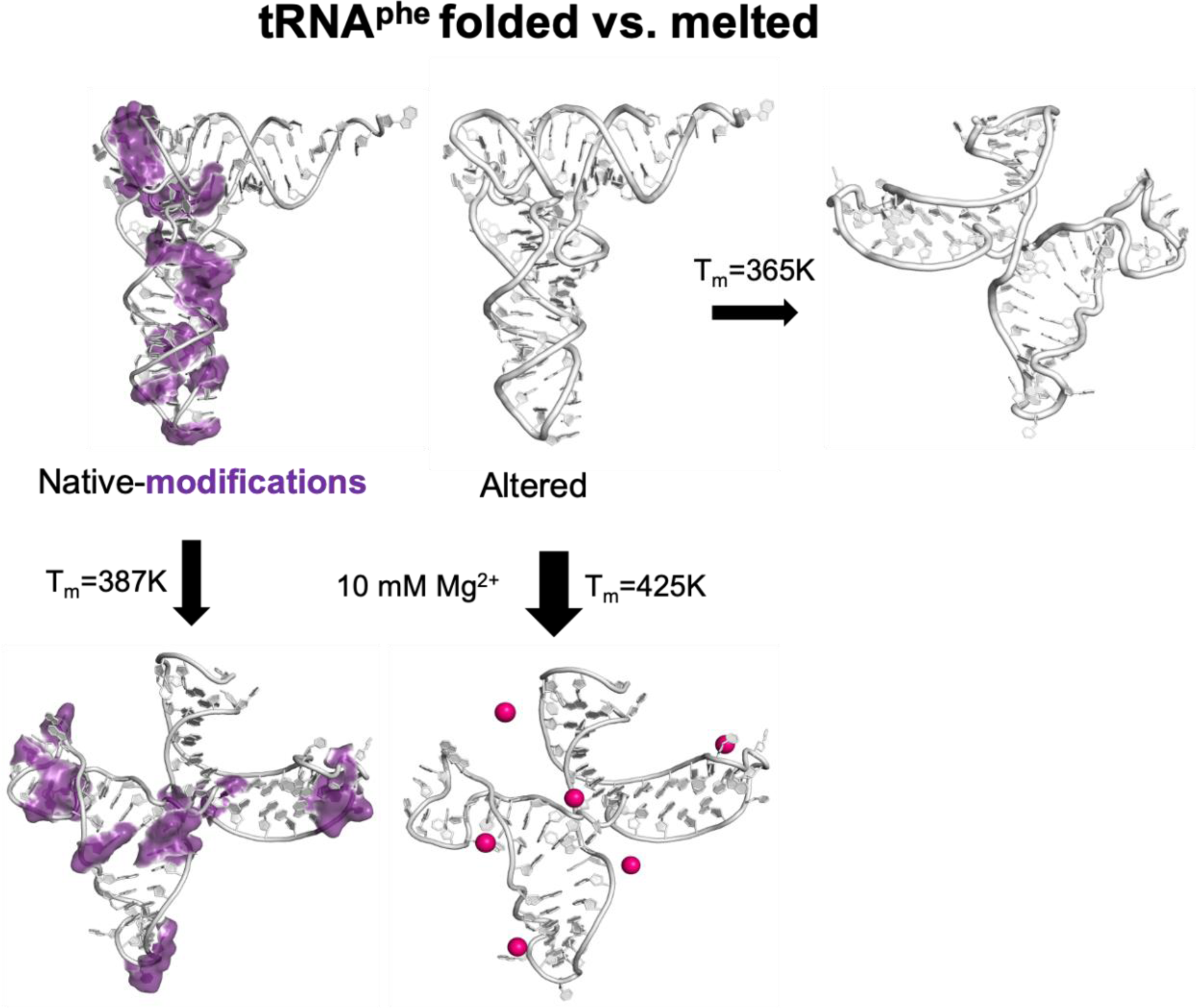

## Introduction

Post-transcriptional modifications, or PTMs, play a critical role in determining the structure and function of RNA, with more than 200 RNA PTMs discovered in various life forms (1). Over 100 of all discovered PTMs are found in transfer RNAs or tRNAs, and tRNAs also have the highest density of modifications, with an average of 13 PTMs found within each tRNA molecule (2, 3). These PTMs affect tRNA stability, codon bias, decoding, and translation efficiency (2).

The tertiary structure of tRNA has been well studied and characterized (4–6). tRNAs have a characteristic cloverleaf secondary structure and an “L” shape when folded, as shown in Figure 1. The tRNA is typically divided into five regions: the acceptor stem, the anticodon loop (ASL), the dihydrouridine (D) loop, the variable loop, and the TΨC loop. One end of the “L” contains the anticodon loop, a critical region for translation, and the opposite end of the molecule presents the amino acid, which is added to a growing polypeptide chain during translation (7). At the elbow or bend of the “L”, the D and TΨC loops interact to support this unique tertiary structure (8).

**Figure 1.**
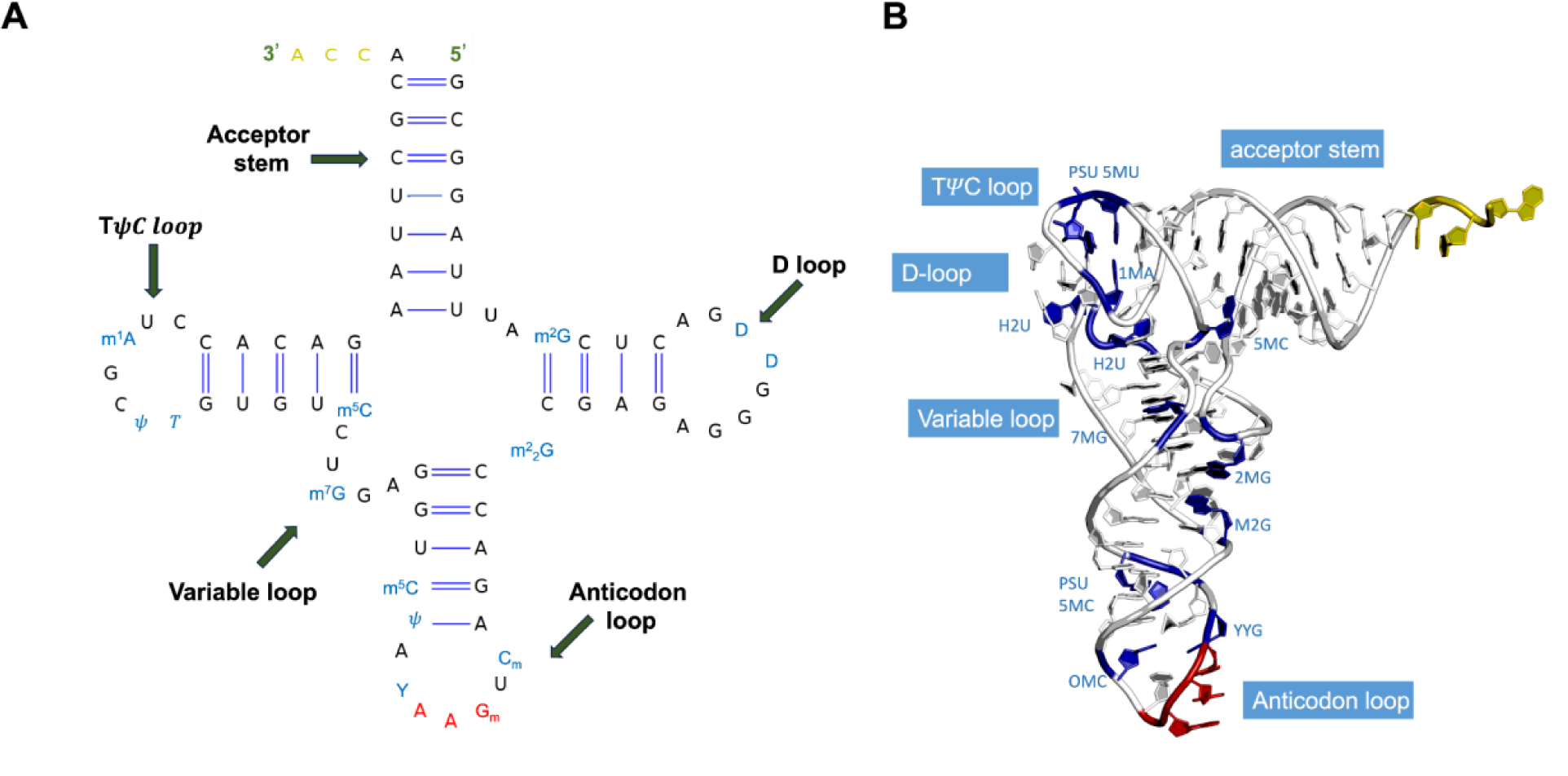
**(A)** tRNA^Phe^ with modifications highlighted in blue and anticodon in red. **(B)** tRNA^Phe^ crystal structure (PDB ID: 1EHZ) cartoon presentation of the tertiary structure; the coloring is consistent with panel A. Secondary structure and tertiary structure were prepared using VARNA: Visualization Applet and PyMOL, respectively (66, 67).

The functions of several PTMs within tRNA remain unknown, but in the last decade, investigations into PTM function and significance have notably increased (9–11). The anticodon loop has been extensively studied, and PTMs within this region have been found to modulate codon-anticodon interaction, regulating translation (12, 13). PTMs within the D loop and TΨC loop have been found to maintain and support the tertiary fold, impacting base stacking (11) and base pairing (14) within the bend.

The role of modifications in tRNA has been previously investigated by targeting a single or a few modifications or by comparing the fully modified and the unmodified tRNA (15). Examples include ASL modifications that affect translation like Queuosine (Q) at position 37 of tRNA^His^ in Escherichia coli (8), 2-thiocytidine (s^2^C), and Inosine (I) on tRNA^Arg^ in Saccharomyces cerevisiae (16), lysidine (L) or granulation at position 34 and 2-methylthio-N6-threonylcarbamoyladenosine (ms^2^t^6^A), on tRNA^Lys^ (17–19), mcm^5^U at position 34 of tRNA^Asp^ in humans (20) and Wybutosine (yW), located at position 37 on tRNA^Gly^ (9) However, structures and biochemical assays of unmodified tRNAs demonstrate that PTMs do not alter the overall L-shaped structure of the tRNA but they are essential to locally orient the nucleotides to maintain functionally active form for various biological processes (21, 22). In addition to current experimental techniques, molecular dynamics simulations have emerged as a powerful tool to reveal the mechanistic details of how these PTMs collectively interact and influence RNA dynamics and structure (23). Here, we chose to study PTMs as an ensemble in tRNA, using a well-established model system tRNA^Phe^ from S. cerevisiae. tRNA^Phe^ from S. cerevisiae has been extensively studied in structural biology to gain insights into modified RNA structure for decades (4, 5, 12, 24–30). Its crystal structure was first determined through X-ray crystallography (4), and NMR studies (5, 26) have provided additional information about the dynamics and conformational changes (25). Besides nucleotide modifications, magnesium (Mg^2+^) has also been shown to be essential for the stability of tRNA’s tertiary structure(27, 31, 32). While there is a wealth of experimentally derived structural information on tRNA^Phe^, the precise functional implications of all its PTMs and the synergy between Mg^2+^ ions and PTMS remain incompletely understood. Theoretical and computational approaches allow for the investigation of each factor’s independent and interdependent effect on tRNA’s conformational stability.

In this study, we use molecular dynamics simulations to examine how the PTMs and Mg^2+^ ions within the native tRNA^Phe^ model contribute to the structural stability of tRNA^Phe^ by comparing a native model containing all of the native PTMS with an altered model, where the PTMs were mutated back to their parent nucleotides (e.g., 1-methyladenosine was mutated back to adenosine and so on) for a range of Mg^2+^ ion concentrations. As tRNA^Phe^ has been studied quite extensively experimentally, we based our hypothesis on their results: that the PTMs within the native model in combination with Mg^2+^ ions would provide prolonged stability when exposed to increasing temperatures within the simulation, resulting in the native model unfolding at a higher temperature range compared to the altered model. We also hypothesized that the two tRNA models would have different unfolding pathways, as previous experimental studies have postulated that a tRNA without PTMs would unfold differently and have other intermediate states compared to a native modified tRNA unfolding under the same temperature range (33).

## Methods

### Model preparation

The structure of tRNA^Phe^ from S. cerevisiae was obtained from the protein data bank (PDB ID: 1EHZ) (4). To investigate the global effects of the modifications within tRNA^Phe^, we prepared a fully altered version, where each modified nucleotide was mutated back to its parent nucleotide (e.g., 7MG is now G in the altered model). The models were edited in MOE (34).

### Parameterizing RNA modifications

There are 11 modified nucleotides at 14 positions found in native yeast tRNA^Phe^, including 2-methylguanosine (m^2^G), 5,6-dihydrouridine (H2U), N2-dimethylguanosine (m2,2G), 2’O-methylcytidine (2’OMC), 2’O-methylguanosine (2’OMG), the Y base or wybutosine (yW), pseudouridine (Ψ), 5-methylcytidine (m^5^C), 7-methylguanosine (m^7^G), 5-methyluridine (m^5^U), and 1-methyladenosine (m^1^A) (Figure 1). We developed Amber-type force-field parameters for each modification. The partial charges were generated using the online RESP charge-fitting server, REDS (35). All modifications were considered with neutral charge except for 7-methylguanosine and 1-methyladenosine, as these two modifications are positively charged in native RNA structures (36, 37). The geometry of each modified nucleoside was energy minimized, and Hartree-Fock level theory and 6–31G* basis sets were employed to arrive at a set of partial charges (38, 39). AMBER-99 force-field parameters were used for bonded interactions, and AMBER-99 parameters with Chen-Garcia corrections were used for Lennard-Jones (LJ) interactions (40, 41).

### Molecular Dynamics Simulation

All-atom molecular dynamics simulations (MDS) were performed using the Gromacs 2019.4 packages (42). Single temperature and replica exchange molecular dynamics (REMD) simulations were performed on both the native (with PTMs) and altered (without PTMs) configurations of the tRNA at varying temperatures and ion concentrations of potassium and magnesium chloride. The Amber-99-Chen-Garcia force field with backbone phosphate modifications (40, 41) was used for nucleic acids. For simulation involving MgCl_2_, we added to the forcefield Mg^2+^ and chloride (Cl^-^) ion parameters developed from the Schwierz lab (43). Trajectories were propagated by integrating Newton’s equations of motion using a Velocity-erlet algorithm with a 2fs time-step. The TIP4P-Ewald (44) model was used to represent water molecules. The temperatures, ion concentrations, and duration of simulations for the different systems have been described below and summarized in the supplemental (Figure S1).

### Single temperature simulations

To study the effect of magnesium on the tRNA structure, we simulated both the native and altered tRNA^Phe^ at five different temperatures (400K, 425K, 450K, 475K, and 500K) and for three different Mg^2+^ ion concentrations (10 mM, 20 mM, and 40 mM) for a total of 15 simulations each. Simulations were run for 200 ns. To add MgCl_2_ to the system, we calculated the number of Mg^2+^ ions to add using the following formula:

num_nmg = (num_wat / 5550) *C_Mg,_

num_wat represents the number of solvent molecules remaining after adding KCl, and C_Mg_ is the concentration of Mg^2+^ ions. We rounded the answer to the nearest integer since fractions of magnesium ions cannot be added to the system. For every Mg^2+^ ion added, we added two chloride ions (num_ncl = num_nmg * 2).

### Replica Exchange Molecular Dynamics Simulations (REMD)

While single temperature simulations are immensely useful in understanding the localized effects of modifications, the sampling is insufficient to capture global dynamics. To investigate melting pathways and the role of modifications and magnesium in maintaining the overall tertiary structure of the tRNA, we performed replica exchange molecular dynamics simulations (45). For the native and altered tRNA^Phe^ models, we first performed all-atom molecular dynamics simulations using a 10.5nm x 10.5nm x 10.5nm cubic box containing the tRNA, 144 K^+^ ions, 70 Cl^-^ ions, and 37217 water molecules, resulting in a concentration of 0.1M KCl. The system was subjected to energy minimization to prevent any overlap of atoms, followed by a 15 ns equilibration run. One dimensional replica exchange molecular dynamics (REMD) simulation with 60 replicas spanning a temperature range of 300K to 440K was performed. The specified minimum swap rate was 0.2, with attempted temperature swaps every 1000 steps (2ps). Coordinates were also saved every 1000 steps. Once equilibrated, the production simulations progressed for roughly 80 ns per replica, resulting in a total of 4.8 μs simulation time for each system.

We also performed REMD simulations on the altered tRNA^Phe^ in MgCl_2_ to understand if Mg^2+^ ions can mimic the effect of modifications on tRNA structure and stability. We only simulated the altered form in a 10mM concentration of MgCl_2_, as this was the highest Mg^2+^ concentration that allowed sampling of both the melted and folded forms of the altered tRNA in our single-temperature simulations. We first performed all-atom molecular dynamics simulations using a 12.45nm x12.45nm x 12.45nm cubic box containing the tRNA, 191 K^+^ ions, 10 Mg^2+^ ions, 20 Cl^-^ ions, and 62687 water molecules. The system was subjected to energy minimization, followed by a 15 ns equilibration run. We performed this REMD simulation with 64 replicas spanning a temperature range between 350K to 450K. Each replica was simulated for 108 ns, resulting in a total effective simulation time of 6.9 μs for the system.

Simulations were analyzed using programs from the Gromacs toolbox to calculate Root Mean Square Deviation (RMSD), Radius of Gyration (Rg), and Root Mean Square Fluctuation (RMSF). The RMSD was calculated for the backbone atoms using the original crystal structure as a reference. Rg was computed about the center of mass in the x, y, and z directions. RMSF was calculated to assess the deviations of each residue’s atomic position from their respective average location within the biomolecular system throughout the simulation.

## Results

### Effect of PTMs and magnesium ions on tRNA^Phe^ structure and stability

We first carried out single temperature MDS on both the altered and native tRNA^Phe^ models at three magnesium concentrations: 10mM, 20mM, and 40mM and at five simulation temperatures: 400K, 425K, 475K, and 500K, as described in the methods. The effect of the selected magnesium concentrations on tRNA^Phe^ structure has been studied experimentally (15, 46). For these single-temperature MDS, we used the RMSD and the RMSF as indicators of the structural integrity and the dynamics of the residues within the models, respectively.

For the native construct, in the presence of 10mM Mg^2+^, we observed that up to 450K, the structure does not deviate significantly from the initial structure, with higher RMSD values observed at temperatures above 475K. In contrast, the altered model shows lowered stability at the same 10 mM Mg^2+^ concentration, sampling higher RMSD conformations at a relatively lower temperature of 425K. This suggests that at this Mg^2+^ concentration, the stability offered by PTMs is not completely compensated by magnesium ions. However, at 20mM Mg^2+^ concentrations, even in the absence of PTMs, the altered model at the different temperatures exhibits behavior similar to the native system, as indicated by comparable RMSD values. At 40mM Mg^2+^ ion concentration, the tRNA maintains its L-shaped structure at the highest tested temperature of 500K. Thus, our simulations demonstrate that the PTMs serve as critical factors in maintaining the structural integrity of tRNA under physiological conditions, working in conjunction with Mg^2+^ ions, as indicated by the results for the altered system at 10mM and 20mM Mg^2+^ ion concentration. At higher concentrations, however, the magnesium ions have the potential to dominate the tRNA structure and stability landscape. These observations are in agreement with previous studies that the loss of PTMs destabilizes tRNA; however, this destabilization can be mitigated by high concentrations of Mg^2+^ (15).

Next, we calculated per-residue root mean square fluctuations (RMSF) to assess the dynamics of individual nucleotides at different temperatures for both native and altered structures at 10 mM Mg^2+^ (Figure 3). While RMSD is indicative of global conformational changes in the overall structure, RMSF provides insights into local dynamics and flexibility. At 400K, the lower end of the explored temperature range, we observe that the RMSF values for the nucleotides in the native and altered tRNA are comparable. However, at 425K and 450K (Figures 3B and 3C), the fluctuations are consistently higher for the altered model compared to the native model, suggestive of higher nucleotide mobility, particularly in the D-loop and the ASL. Both these loops are structurally and functionally important. While the conformation and composition of the ASL plays a crucial role in aminoacylation, ribosome binding, and translation, the interaction of the D-loop with the TΨC loop is critical in maintaining the tertiary structure of the tRNA. Based on the RMSF values at 425K, we observe a strong deviation of the altered from the native constructs at four sites - dihydrouridine (D) at positions 16 and 17 in the D-loop and OMC, and OMG modifications at positions 32 and 34 in the ASL (Figure 3B, red circles). Five of the fourteen modified nucleotides in tRNA^Phe^ are clustered around the interacting D-loop and TΨC loops. Besides the canonical G_19_:C_56_ base pair, a network of inter-loop and intra-loop interactions rely on modified nucleotides to stabilize the region, including polar contacts between D_16_:U_59_, G_18_:Ψ_55,_ and m^5^U_54_:m^1^A_58_ and stacked nucleotides G_57_, m^1^A_58,_ and G_18_. We observe that in the simulated native construct with 10mM Mg^2+^, at 400K, most of these interactions are maintained, consequently conserving the D-loop and TΨC loop interactions. However, in the altered state, at the same temperature and Mg^2+^ ion concentration, our simulations show a loss of most of these interactions, indicating the importance of modifications in orienting the nucleotides to establish the necessary contacts for this kissing-loop interaction (Figure 4). For the same 10 mM Mg^2+^ concentrations, at higher temperatures, the reduced stability in the altered model becomes increasingly pronounced (Figure 3), giving greater weight to the hypothesis that PTMs provide structural stability in key regions in tRNAs.

**Figure 2.**
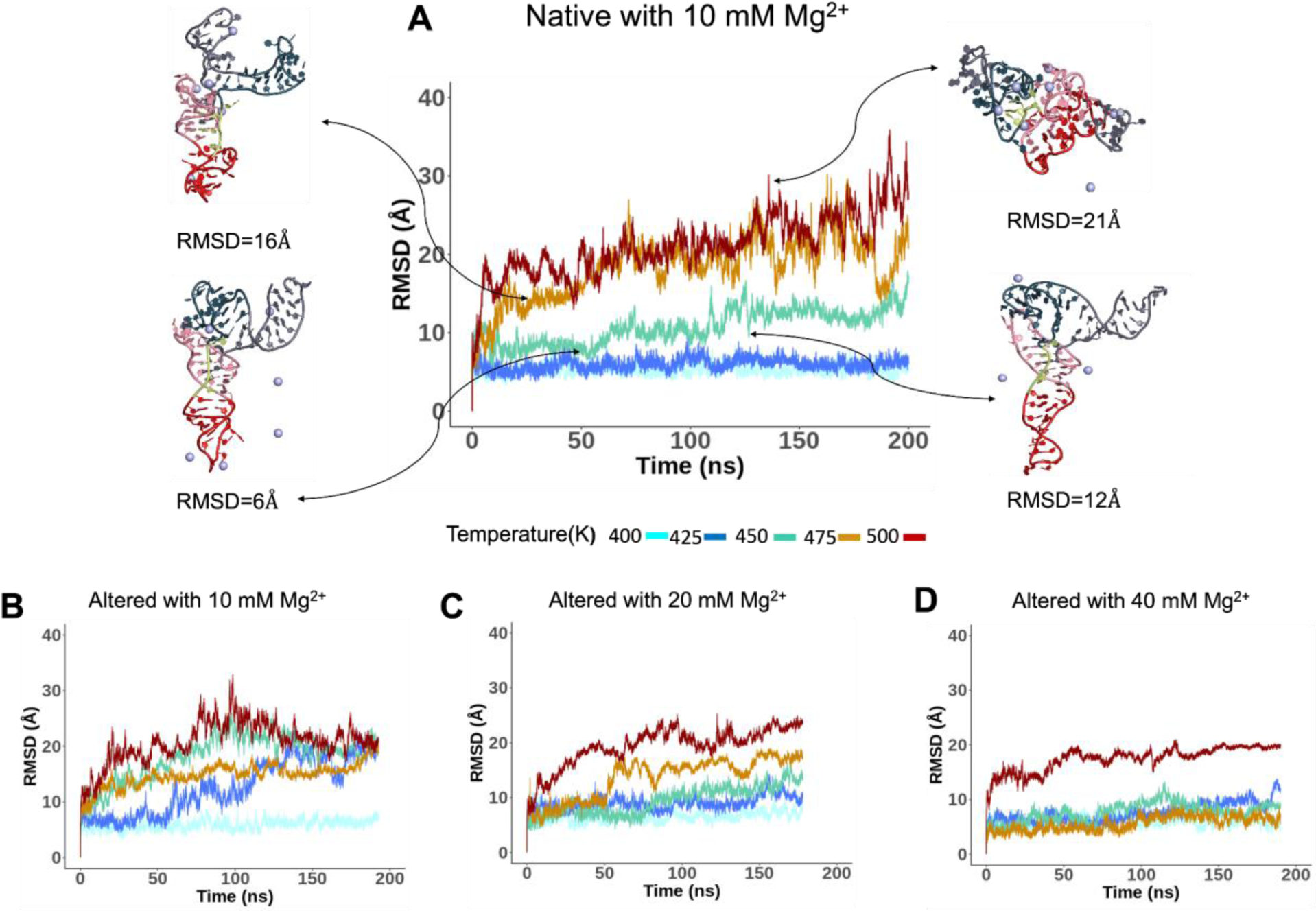
**(A)** RMSD and representative snapshots of Native tRNA^Phe^ temperatures (400K, 425K, 450K, 475K, and 500K) at 10 mM Mg^2+^. RMSD of altered tRNA^Phe^ at **(B)** 10 mM Mg^2+,^ **(C)** 20 mM Mg^2+,^ **(D)** 40 mM Mg^2+^.

**Figure 3.**
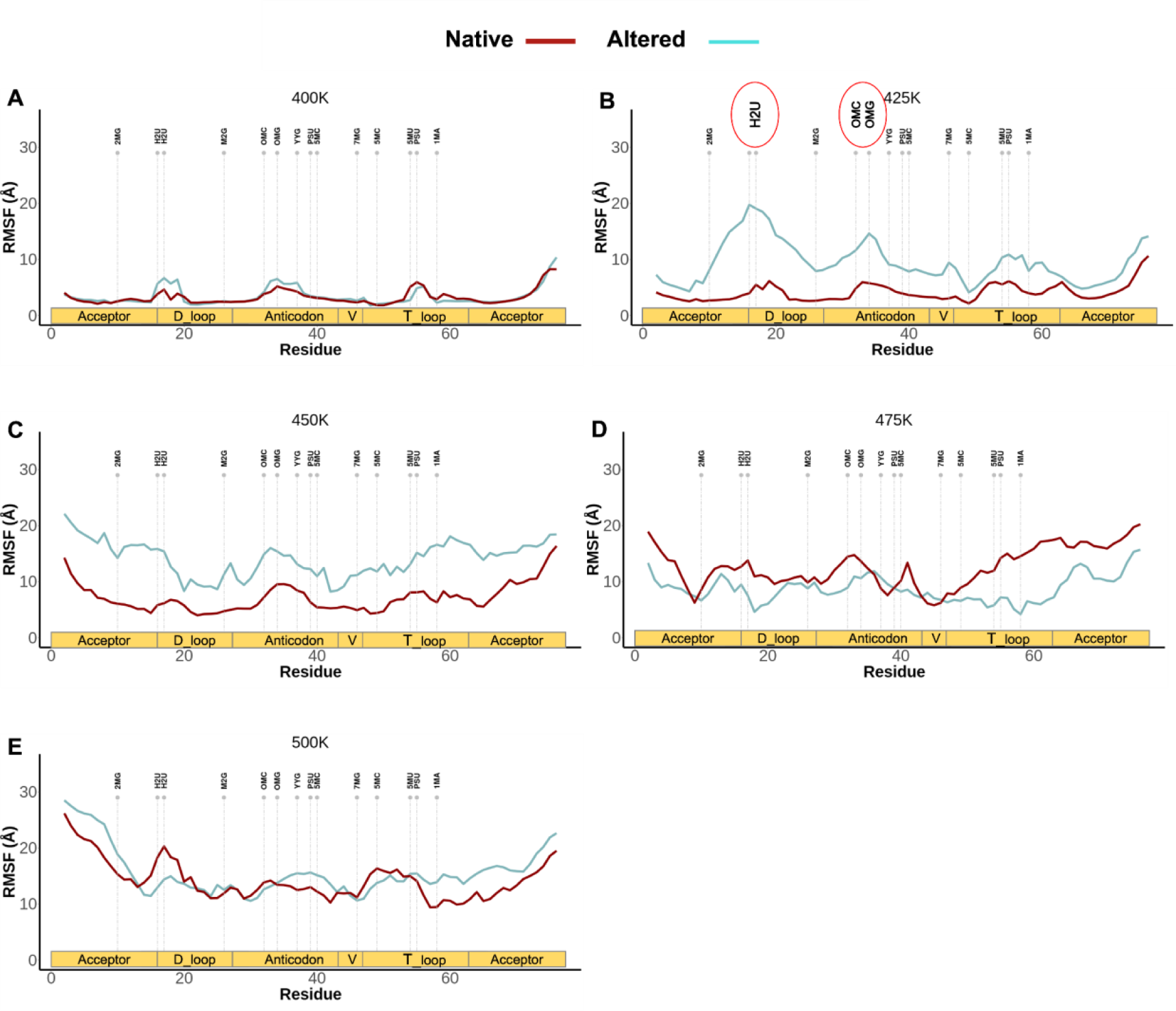
RMSF plots for the native and altered structures in 10 mM Mg^2+^ at temperatures **(A)**400K, **(B)**425K,**(C)** 450K,**(D)** 475K, and **(E)** 500K. The vertical dash lines indicate the locations of modifications.

**Figure 4.**
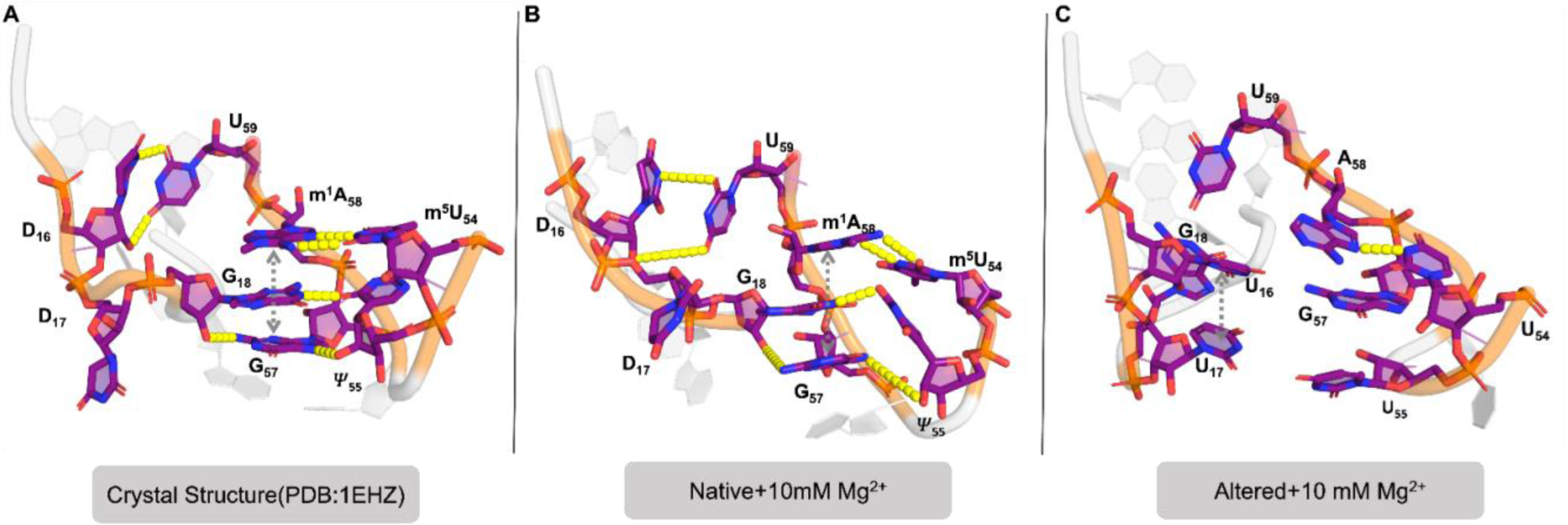
D-loop - TΨC loop interaction observed in **(A)** crystal structure (PDB: 1EHZ), **(B)** simulated native tRNA structure at 400K and 10mM Mg^2+^, and **(C)** simulated altered tRNA structure at 400K and 10mM Mg^2+^. The simulated structures are representative of the dominant configuration observed during the simulation.

### Effects of PTMs and Mg^2+^ on tRNA^Phe^ folding dynamics

Our analysis thus far shows that PTMs and Mg^2+^ play a critical role in stabilizing the overall tertiary structure of the tRNA. Next, we asked how PTMs and Mg^2+^ affect the melting pathway of the tRNA. To address this, we conducted REMD simulations on both the native and altered models of tRNA^Phe^, as explained in the methods section. For each construct, 60 replicas were generated and simulated in parallel at temperatures ranging from 300K to 450K.

In order to understand melting, we explored different metrics to find a variable that best differentiated the folded and unfolded states. We first calculated the radius of gyration (Rg) and the RMSD of the tRNA for each frame in the REMD simulations using the original crystal structure as a reference, as described in the methods. The population density of RMSD, Rg pair is illustrated utilizing a contour heat map to identify highly populated areas, indicating possible intermediates along the unfolding pathway. Figure 5 shows the representative heat maps of the native and altered tRNA^Phe^ models with the structures of high population areas within the heat map alongside.

**Figure 5.**
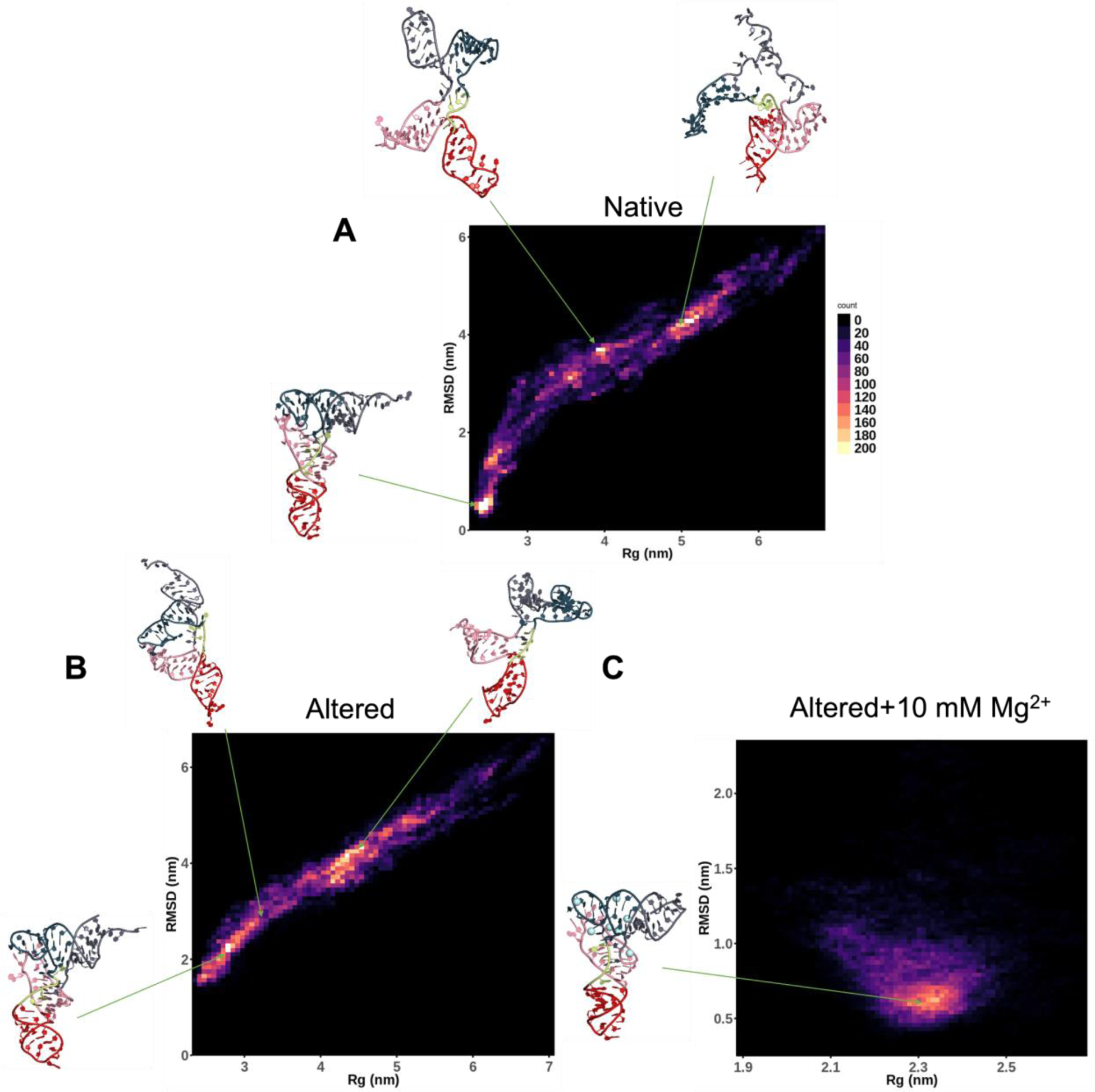
RMSD vs. Rg heatmap for **(A)** native **(B)** Altered **(C)** Altered+10 mM Mg^2+^ tRNA^Phe^, where RMSD is on the y-axis, and Rg is on the x-axis. Both RMSD and Rg are in nm. This heatmap is based on one of the replica simulations produced as a result of the REMD simulations. The structures representing the indicated areas of the high population within the heat map have been included.

The most highly populated structure of the native tRNA^Phe^ unfolding pathway is closest to the crystal structure, as shown in Figure 5A. The stem loops are still structured, and the overall L-shape is maintained via the kissing loop interaction between the D-loop and the TΨC-loop. As the simulation progresses, this kissing loop interaction is lost; the stem loops extend outward, resulting in a structure like the t-shaped secondary structure map (Figure 1A). Eventually, the interior loop bulges out and expands, slowly but surely disrupting the stem loops’ secondary structure, leading to a structural population with the highest RMSD and Rg values.

The altered model, however, demonstrates a different unfolding pathway. Overall, the high population areas are more diffuse and less defined compared to the native construct. This suggests that the PTMs in the native construct actively prevent the loss of the tRNA structure compared to the altered model’s unfolding pathway (Figure 5B). As shown by representative structures, at a lower RMSD and Rg, the internal bulge by the kissing loop interaction between the D-loop and TΨC-loop is wider than in the native construct. The bulge widens, and the anticodon stem loop structure becomes disordered as the structure unfolds. While both the native and the altered structures adopt the t-shaped structure, as observed by high density regions around RMSD and Rg at 4nm, they demonstrate different propensities to sample the conformation, as indicated by the well-defined vs. distributed high-density region in the heatmaps of the native and altered constructs respectively.

Mg^2+^ significantly affects the tRNA structure, confining it to a more compact and stable conformation in an altered form. At 10mM Mg^2+,^ the heat map shows a much smaller range for RMSD and Rg sampled than the native and altered construct without magnesium (Figure 5C). The heat map reveals a distinct circular pattern and increased sampling in regions with RMSD less than 2 nm, indicating a native like structure is maintained throughout the simulations. This observation suggests that the presence of Mg^2+^ offsets the lack of PTMs and stabilizes the tRNA^Phe^ structure by restricting conformational fluctuations and promoting a more rigid conformation.

To gain insights into the conservation of local structure, we calculated the RMSF for each residue within the native and altered models across all replicas (Figure 6). The fluctuations of the stem nucleotides are lower than those in the loop nucleotides, and this trend is observed even at higher temperatures, although it is less remarkable. This indicates that the secondary structure is initially retained while tertiary contacts are lost as the tRNA begins to melt. The presence of Mg^2+^ ions reduce the overall flexibility of the nucleotides even at higher temperatures.

**Figure 6.**
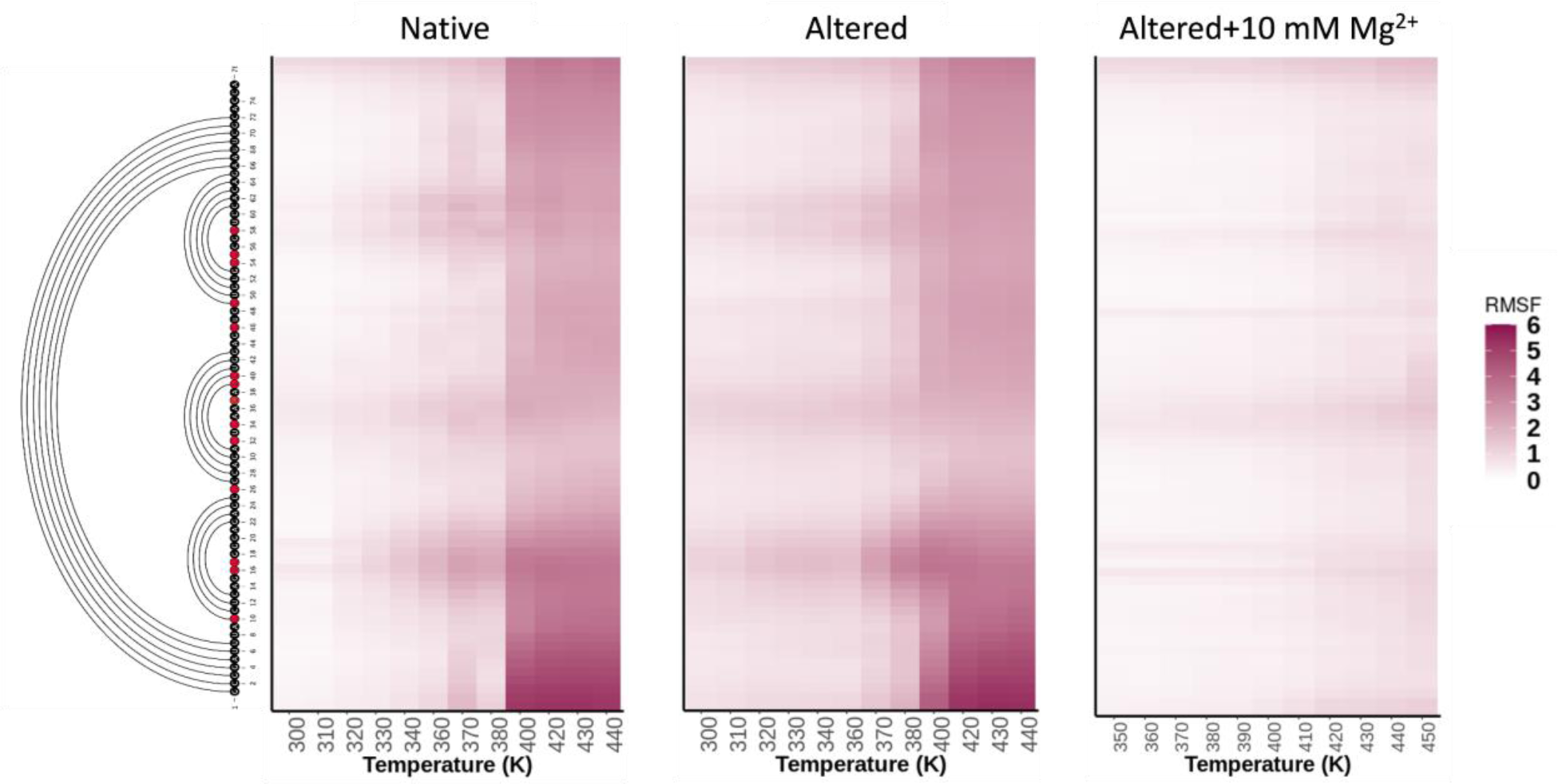
RMSF of the native, altered tRNA^Phe,^ and altered tRNA^Phe^ +Mg^2+^(10mM) systems at different temperatures.

In addition to the melting pathways, we also generated simulated melting curves. Using the heatmap analysis as a reference, we used the kissing loop interaction between the D-loop and the TΨC-loop to classify the tRNA structures obtained from the REMD simulations into “folded” and “unfolded” states. The distance between G19:C56 in the kissing loop with the same cutoff of 15Å as this metric best distinguished the unfolded states from the folded ones (Figure S2). The fraction of folded structures was then calculated as a function of the simulation temperature for each case. Consequently, a “melting” curve was plotted, like the traditional experimental UV melting trends (Figure 7).

**Figure 7.**
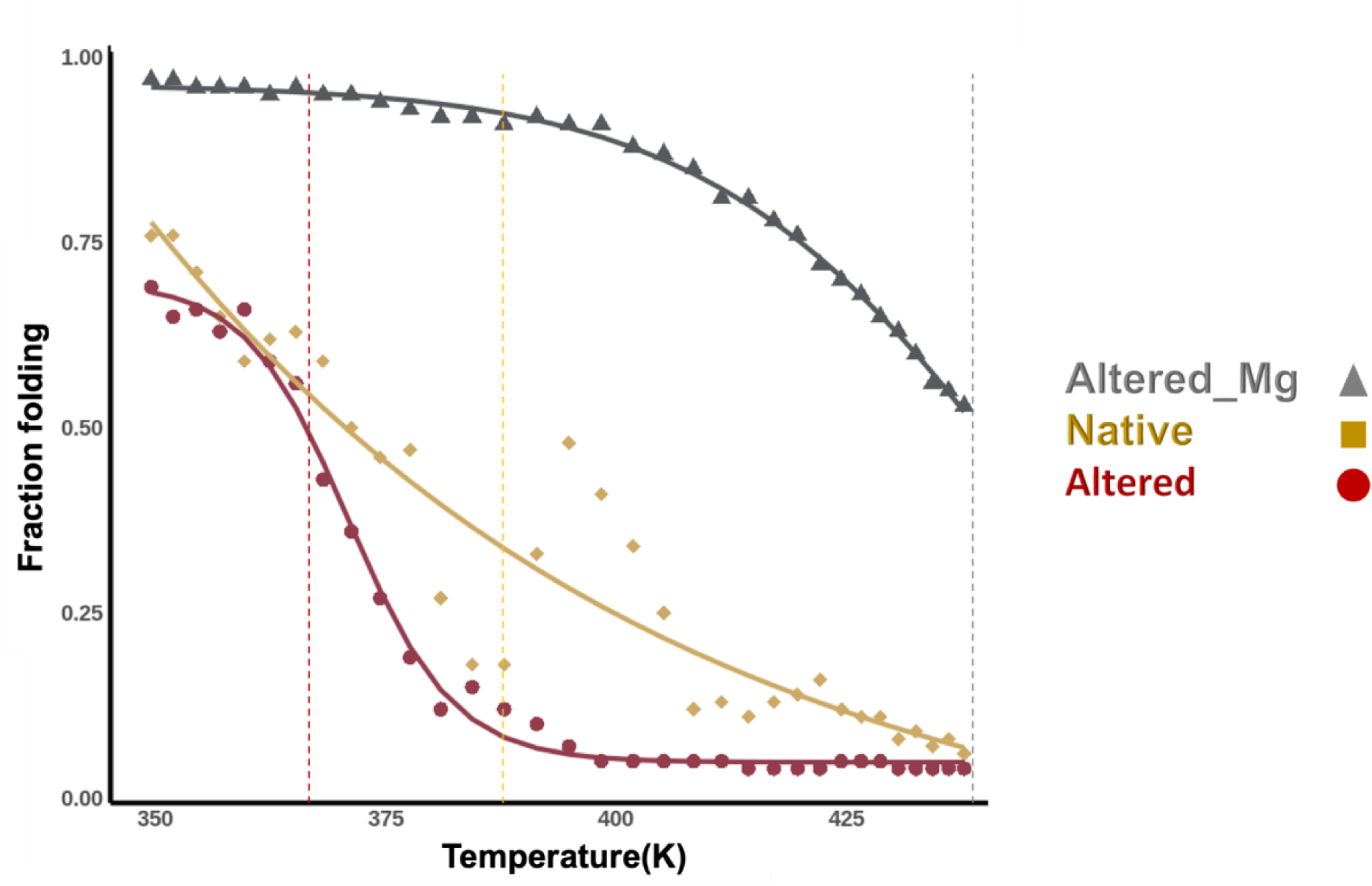
Folded vs. Simulation Temperature for all the native and altered tRNA^Phe^ model, and altered tRNA^Phe^ model with 10mM Mg^2+^.

Extending the UV melting analysis to the melting curves obtained from our simulations, the temperature at 50% of the structures folded can be interpreted as the “melting temperature.” The native tRNA maintains a more “folded” state overall and hits 50% folded around 387K compared to the altered model, reaching a 50% folded state around 365K. This difference in “melting” temperature reaffirms that the altered model loses its structure much earlier in the temperature range than the native model. Interestingly, with the addition of 10mM Mg^2+^, the altered model reaches a 50% folded state at a much higher temperature of ∼450K, thus indicating that high concentrations of Mg^2+^ can match and even enhance the stability imparted by the PTMs on tRNA structure.

### Interplay between PTMs and Mg^2+^ binding sites in tRNA^Phe^

Our analysis has established that both PTMs and Mg impart structural stability to tRNA^Phe^, and their relative contributions to stability are concentration dependent. Furthermore, high enough Mg^2+^ concentrations even compensate for stability loss due to the absence of PTMs. We further investigated the spatial distribution of Mg^2+^ ions in the vicinity of the native and altered tRNA^Phe^ at 400K to find links between PTMs and Mg^2+^ binding. The choice of 400K as the reference temperature was driven by its consistently low RMSD values across tRNA^Phe^ structures, indicating stability and native conformation. This ensured an ideal baseline for studying Mg^2+^ ion behavior. The spatial distribution of Mg^2+^ ions within 3.3 to 10 Å of each nucleotide in both the altered and native tRNA models was mapped onto the structure as a density map (Figure 8), with darker shades representing a higher frequency of finding Mg^2+^ ions in the vicinity of the nucleotide during 200ns MDS. In the native tRNA structure, at 10mM Mg^2+^ concentration, the ion distribution is consistent with the binding sites observed in the crystal structure (Figure S3). This alignment with the crystal structure reaffirms the specificity and importance of these Mg^2+^ ion binding sites. For the altered tRNA, at the same 10mM Mg^2+^ concentration, fewer Mg^2+^ ions bind in the anticodon loop region (Figure 8B), while more Mg^2+^ are observed around the V-loop junction. At the D-loop: TΨC-loop interaction site, Mg^2+^ binding is consistent in both the native and altered constructs (Figure 8A, B). These observations reveal that while the crystal structures indicate that binding of Mg^2+^ to tRNA is mainly via backbone interactions and largely non-specific, the PTMs play a critical role in creating Mg^2+^ binding sites by inducing local specific 3D conformations (Figure 9B) (15). At higher Mg2+ concentrations, the Mg^2+^ density map shows a significant increase of Mg^2+^ ions throughout the tRNA in its altered state (Figure 8C, 8D), with several of these binding sites situated in regions with high PTMs. Thus, at elevated Mg^2+^ concentrations, Mg^2+^ ions emerge as dominant factors shaping the tRNA structure, compensating for the lack of PTMs and reinforcing the overall structural integrity (Supplemental Video SV1 showing the unfolding of tRNA^Phe^ in its native and altered state at 425K and 10mM Mg^2+^).

**Figure 8.**
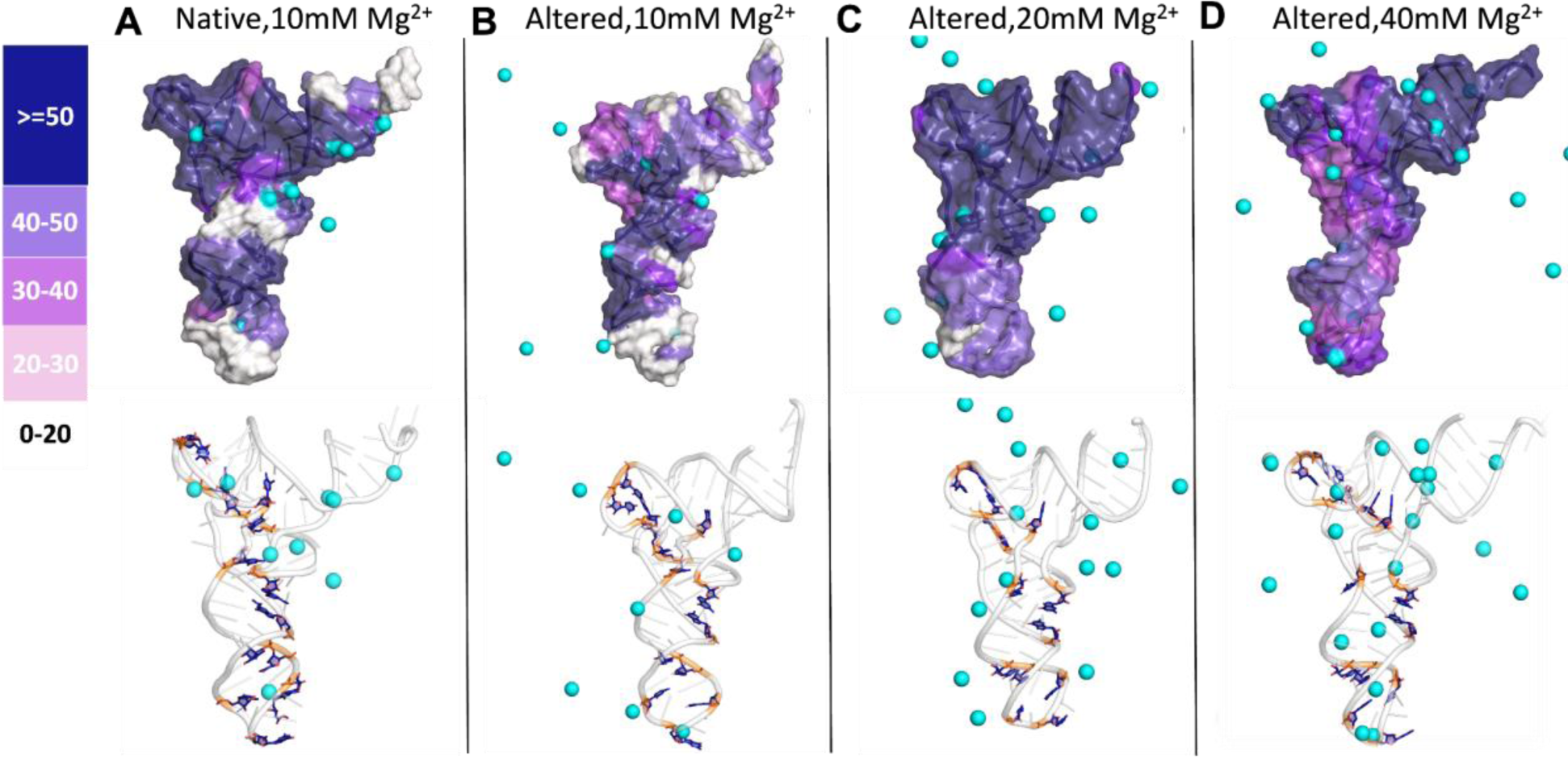
Spatial distribution of Mg^2+^ ions around altered and native models at varying Mg^2+^ concentrations at temp 400K. The top figures in each column depict color intensity in the heatmap, ranging from dark blue for the highest number of residents to white for the lowest number, with pink and purple shades indicating intermediate status. In the second row, the cartoon representation illustrates the same tRNA^Phe^ as in the first row, with cyan spheres depicting Mg^2+^ ions surrounding the structure and dark blue residues signifying modified residues. All the structures were captured from clusters with the highest number of snapshots during the simulation.

**Figure 9.**
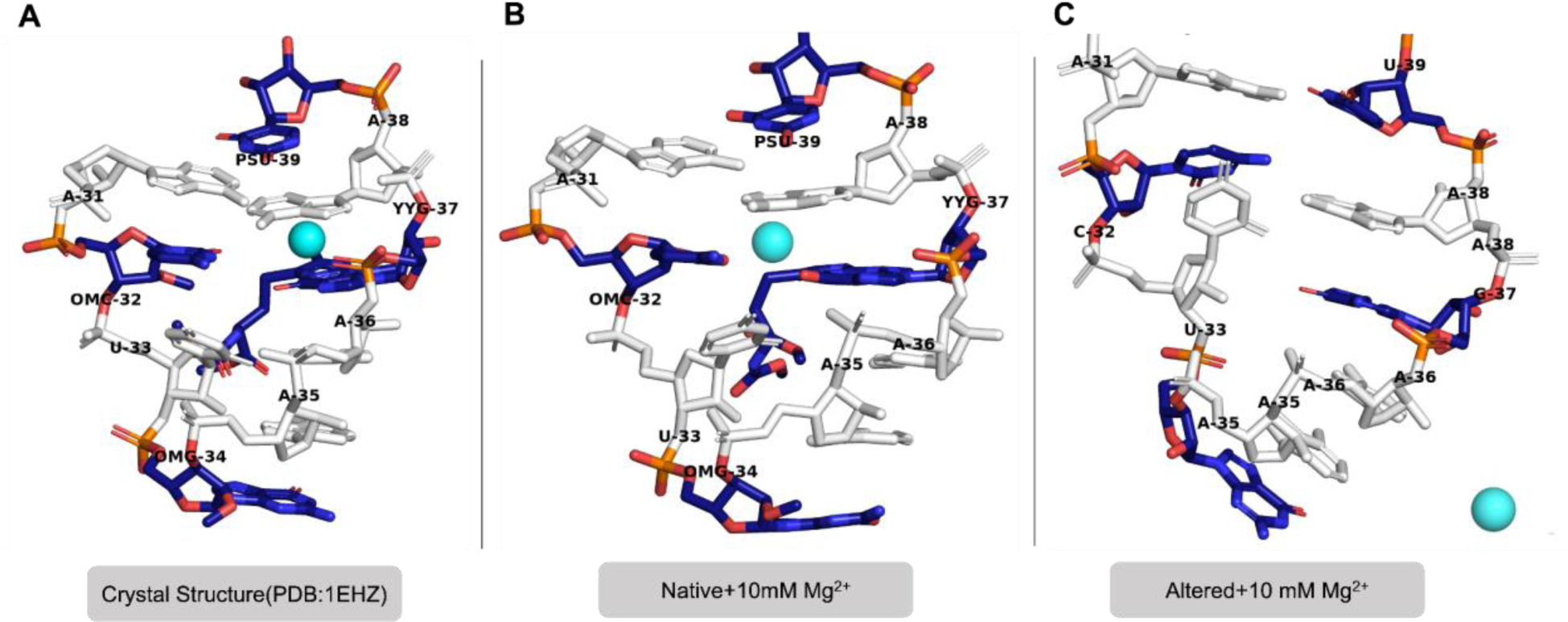
Comparative analysis of the anticodon loop region within tRNA^Phe^ under 10 mM Mg^2+^ concentrations, featuring the **(A)** crystal structure (PDB: 1EHZ), **(B)** simulated native tRNA structure, and **(C)** simulated altered tRNA structure. Modifications are labeled and colored in blue. Mg^2+^ ions are represented as cyan spheres. The second and third panels were captured from clusters with the most snapshots during the simulation.

### Discussions

In this study, we investigated the effects of PTMs and Mg^2+^ ions on the structure and stability of tRNA^Phe^. Our findings shed light on the intricate interplay between these factors and provide insights into the molecular mechanisms governing tRNA folding energy landscapes. While Mg^2+^ ions are primarily linked with RNA folding and stability, and PTMs are typically explored for their functional roles, our simulations suggest that it is the synergy between PTMs and Mg^2+^ ions that results in the maintenance of the tRNA structure. Specifically, we showed that PTMs can influence the binding of Mg^2+^ ions by local rearrangements in the different segments of the tRNA.

It has been shown that PTMs are critical to the structure and stability of tRNAs (47, 48). In agreement with these previously published results, we find that when PTMs are removed, such as in our altered model, there is a noticeable effect on the conformations and stability of the folded tRNA. For example, at the junction of the D-loop and TΨC loop, the absence of modifications renders these segments of the altered tRNA significantly more flexible compared to the native tRNA, as indicated by the higher RMSFs even in the presence of 10 mM Mg^2+^ (Figure 3B). On the TΨC loop, Ψ_55_ is critical to maintaining the U-turn configuration via direct and water-mediated hydrogen bonding with neighboring nucleotides (49), as has been previously observed. We speculate that the replaced U_55_ disrupts this U-turn, affecting the orientation of the downstream nucleotides at positions 57 and 58, which in turn interrupts the cross-stacking interactions between the D-loop and the TΨC loop. On the other hand, in the native D-loop, the non-planar dihydrouridines prevent stacking, thus imparting more flexibility to the loop and promoting the interactions of D_16_ with U_59_ and G_18_ with the TΨC loop nucleotides. When replaced with uridines, in the altered state, we observe a stacked conformation between U16 and U17 and no consistent interactions with the neighboring nucleotides, which destabilizes the whole region. This result aligns with and complements experimental findings, which demonstrated that the modified D-arm, with dihydrouridine, folded into a stable hairpin, while its unmodified counterpart exhibited dynamic, interconverting conformations—underscoring the importance of the modifications in maintaining and supporting the tRNA D-loop (50, 51). The disruption of the D_16_: U_59_ interaction (Figure S6) is also observed in the experimentally solved structure of the unmodified structure of tRNA^Phe^ in *E. coli (50)*.

The role of magnesium ions in tRNA stability has been extensively explored. Early work by Stein and Crothers (1976) and Rhodes (1977) highlighted the significance of magnesium in promoting tRNA structural stability and its contribution to the thermal unfolding process (52, 53). Our study, in line with these observations, underscores the stabilizing influence of magnesium ions on tRNA structure. We observed that at high Mg^2+^ concentrations (20mM and 40mM), the tRNA with no modifications maintains its native L-shaped structure even at elevated temperatures. This aligns with the previous notion that magnesium ions play a central role in stabilizing tRNA’s secondary and tertiary structure (15, 54, 55).

In tandem with our computational analyses, we draw upon the wealth of information from previous crystallographic studies (56–60). Notably, the studies consistently report the presence of magnesium ions in crucial regions, including the kissing loop, anticodon loop, and D loop. The findings from crystallography align closely with our computational results, revealing a shared recognition of the importance of Mg^2+^ binding sites in stabilizing tRNA structures(Figure S2).

To corroborate our computational observation, we turn to the work of Friedrich and Hagermann (1997), Leroy et al. (1977), Maglott et al. (1998), and Maglott et al. (1999) on yeast tRNA^Phe^ (48, 61–64). These experimental studies on yeast tRNA^Phe^ emphasize the crucial role of Mg^2+^ in stabilizing tRNA structures. Specifically, these investigations highlight that the folding of key regions in tRNA, such as the 8–12 turn and the T-loop/D-loop core, strongly depends on Mg^2+^ concentration. Our simulations support these findings and, more interestingly, suggest a synergy between PTMs and Mg^2+^ ions in maintaining the tRNA structure. We showed that PTMs can influence the binding of magnesium ions by local rearrangements in the different segments of the tRNA. For instance, we observed contrasting effects on binding a single Mg^2+^ ion between the native and altered structures due to variations in stem-loop conformations (Figure 9B, C). Our findings reveal that PTMs enhance tRNA stability at 10 mM Mg^2+^ ion concentration.

Furthermore, our REMD simulations enabled us to explore conformational dynamics and unfolding pathways in unprecedented detail. Using melting curve analysis inspired by Kirk et al. (2008), we quantified the folding transitions of the native and altered tRNA models, demonstrating the stability-enhancing effects of PTMs and magnesium ions (65).

In conclusion, this study advances the understanding of tRNA stability by investigating the interplay between PTMs and magnesium ions. We provide a nuanced view of tRNA folding energy landscapes by integrating multiple analytical techniques and comparing our results to pertinent literature. Overall, this work is crucial in expanding our knowledge of tRNA folding dynamics.

## Data Availability

The data and molecular models used and analyzed during the present study are available from the corresponding author on request.

## Acknowledgments

We would like to thank Dr. Christopher Myers and Isabel Dengos for their help with scripts and code for plot generation.

